# Genome-Wide Linkage Disequilibrium, Haplotype Block Structure, and Population Diversity in Nili-Ravi Buffalo (*Bubalus bubalis*)

**DOI:** 10.64898/2026.02.22.707281

**Authors:** Atiq Ahmad, Hamid Mustafa, Waqas Ahmad Khan, Abdul Manan, Irem Anwer, Waqas Akram

## Abstract

Linkage disequilibrium (LD) and haplotype block structure govern the resolution and utility of genomic selection, marker-assisted selection, and genome-wide association studies (GWAS) in livestock. We performed a comprehensive genome-wide characterization of LD decay, haplotype block architecture, and population diversity across all 24 autosomes in Nili-Ravi buffalo (*Bubalus bubalis*; n = 85), using 43,543 post-quality-control SNPs. Mean genome-wide r^2^ was 0.124 (median 0.074) and mean D’ was 0.540 (median 0.481), with LD half-decay at ≈70 kb. A total of 133 haplotype blocks encompassing 721 SNPs were identified (Gabriel et al., 2002). Haploview analysis of nine chromosomes harbouring bTB resistance candidate genes revealed contrasting selection signatures: directional selection at innate immune loci (IFNG, TLR1; H’ < 0.55) versus balancing selection at adaptive immune loci (BoLA-DRB3, SP110; H’ > 1.0). Critically, BBU15 Block 3 (28.6 kb; OR52E5/NCR1 locus, 47.16 Mb) showed a genome-wide significant integrated haplotype score (iHS; –log_1 0_ p = 5.408), directly co-localising with the published bTB susceptibility QTL (Bermingham et al., 2014). The TAA haplotype (frequency 53.3%) at this block represents a candidate resistance-associated haplotype for marker-assisted selection. These findings provide essential parameters for SNP panel design and bTB resistance breeding in South Asian buffalo.

## INTRODUCTION

The Nili-Ravi buffalo (*Bubalus bubalis*) is the most economically important water buffalo breed in Pakistan, contributing over 60% of national milk production (FAO, 2023). Despite its significance, the genomic architecture of this breed, particularly the extent and structure of linkage disequilibrium (LD), remains poorly characterized compared to taurine dairy cattle. Understanding LD structure is foundational for GWAS power, genomic selection reference panel design, and marker-assisted selection for complex traits including bovine tuberculosis (bTB) resistance (Slatkin, 2008; Meuwissen et al., 2001). Bovine tuberculosis, caused by *Mycobacterium bovis*, poses a severe economic and public health burden in Pakistan, with buffalo serving as a major reservoir (Javed et al., 2010). Identification of haplotype blocks at bTB resistance loci provides a foundation for genetic selection programmes to improve population-level disease resistance.

Two LD metrics are employed throughout: r^2^, the squared allelic correlation quantifying the proportion of variance explained between loci; and D’, Lewontin’s normalized coefficient sensitive to historical recombination. Haplotype block detection using the Gabriel et al. (2002) confidence interval method partitions chromosomes into discrete segments of suppressed recombination, within which allele combinations are highly predictable. The Haploview software (Barrett et al., 2005) provides graphical visualization of these blocks as colour-coded triangular LD matrices where red indicates D’ = 1.0 (complete LD) and progressively lighter shading indicates diminishing LD. This study characterizes genome-wide LD decay, haplotype block architecture, and specifically examines Haploview LD patterns at nine chromosomes harbouring bTB resistance candidate genes in Nili-Ravi buffalo.

## MATERIALS AND METHODS

### ETHICAL STATEMENT

This study used previously generated genotype data obtained from published sources and a public repository. No new biological sampling, animal handling, or experimental interventions were conducted for the present analyses. Ethical approvals for the original sample collection were secured in the respective primary studies and/or institutional frameworks under which the datasets were generated. Public data were accessed and used in accordance with repository terms and conditions; therefore, additional ethical approval was not required for this secondary analysis.

#### Genotype Datasets and Populations

Genome-wide SNP genotypes generated using the Axiom 90K buffalo array were assembled from three independent sources. First, genotypes for 35 buffalo were obtained from the HEC-NRPU (No. 16844) funded project. Second, genotypes for 35 animals were incorporated from a previously reported dataset (Irem, 2022). Third, publicly available genotypes for 15 Nili-Ravi and 10 Kundi buffalo were downloaded from Harvard Dataverse (2024; https://doi.org/10.7910/DVN/UGA2QX).

Before downstream analyses, datasets were harmonized for SNP identifiers, chromosome labels, genomic coordinates, and allele coding. Variants with unresolved allele/strand inconsistencies across datasets were excluded to avoid spurious ROH calls or inflated differentiation signals.

#### Quality Control and Marker Filtering

Quality control (QC) was applied at the marker level to retain high-confidence SNPs for ROH and selection analyses. SNPs were removed if they met any of the following criteria: (i) minor allele frequency (MAF) < 0.05, (ii) genotype call rate < 0.90, or (iii) significant deviation from Hardy–Weinberg equilibrium (p < 0.001). Only autosomal SNPs passing QC were retained, and sex chromosomes were excluded to prevent sex-linked marker behavior and hemizygosity from biasing homozygosity-based metrics. After QC, 43,389 autosomal SNPs were available for analyses.

#### SNP Density Profiling

To evaluate marker coverage across the genome, SNP density was summarized using non-overlapping 1-Mb autosomal windows. The number of retained SNPs per window was calculated and visualized as a genome-wide heatmap. This step provided a diagnostic view of marker distribution and identified chromosomal segments with sparse or dense SNP coverage that could influence ROH detection power and regional signal interpretation.

#### Linkage Disequilibrium and Haplotype Blocks

Pairwise r^2^ and D’ were computed for all SNP pairs within 500 kb using PLINK v1.90 (--r2, --ld-window-kb 500). Values were binned into seven distance categories (<30, 31–70, 71–100, 101–200, 201–300, 301–400, 401–500 kb). Haplotype blocks were defined by the Gabriel et al. (2002) confidence interval method (PLINK --blocks). Haploview v4.2 (Barrett et al., 2005) was used to generate LD matrix plots and estimate haplotype frequencies for each block.

#### Selection Signature and bTB Candidate Gene Analysis

The integrated haplotype score (iHS; Voight et al., 2006) was computed using the rehh package v3.2.1 in R after phasing with SHAPEIT v2. Thirteen bTB resistance candidate gene loci were annotated across 9 chromosomes. For each candidate chromosome, Haploview LD plots were examined to identify blocks co-localising with candidate gene coordinates. Shannon diversity (H’ = −Σpiln pi) was calculated per block as a quantitative diversity index.

## RESULTS

### SNP Panel and Genome-Wide LD Decay

After QC, 43,543 SNPs were retained across 24 autosomes (Table 1). Mean r^2^ declined from 0.31 at <30 kb to 0.09 at 401–500 kb (LD half-decay ≈70 kb; Figure 1). Mean D’ remained elevated (0.52–0.78) throughout the 0–500 kb range (Figure 1, dashed). The r^2^ scatter plot (Figure 2) confirms wide LD variation with LOESS mean at r^2^ ≈ 0.37 at 0 kb declining to 0.12 at 500 kb. Per-chromosome D’ was remarkably uniform (range 0.524–0.558; Figure 3), while r^2^ showed greater inter-chromosomal heterogeneity (Figure 4). A total of 133 haplotype blocks (mean 225.09 kb; range 20.71–1,754.53 kb) were identified genome-wide (Table 3).

**Figure 1.**
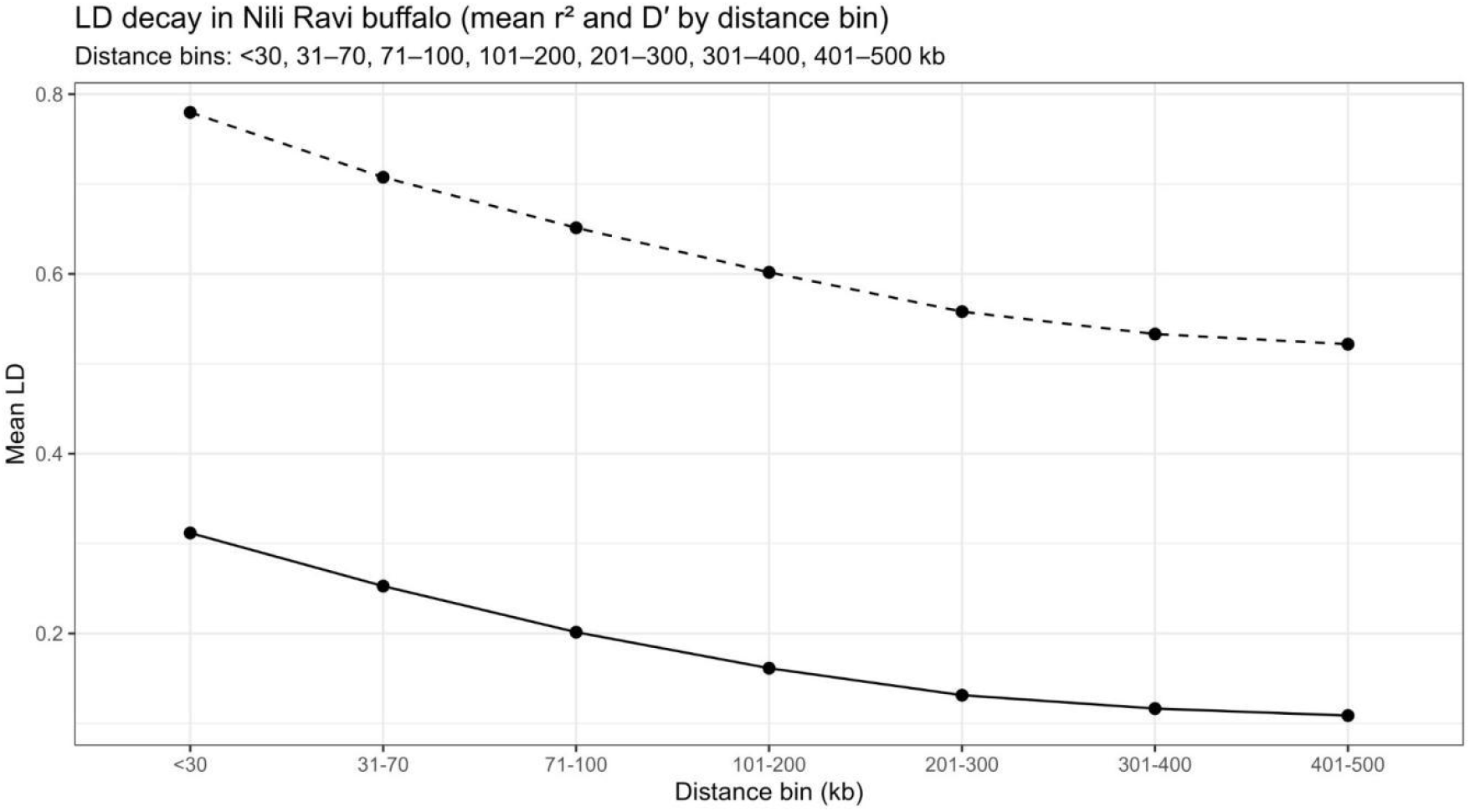
Genome-wide LD decay in Nili-Ravi buffalo. Mean r^2^ (solid) and D’ (dashed) across seven distance bins. The half-decay distance for r^2^ is approximately 70 kb.

**Figure 2.**
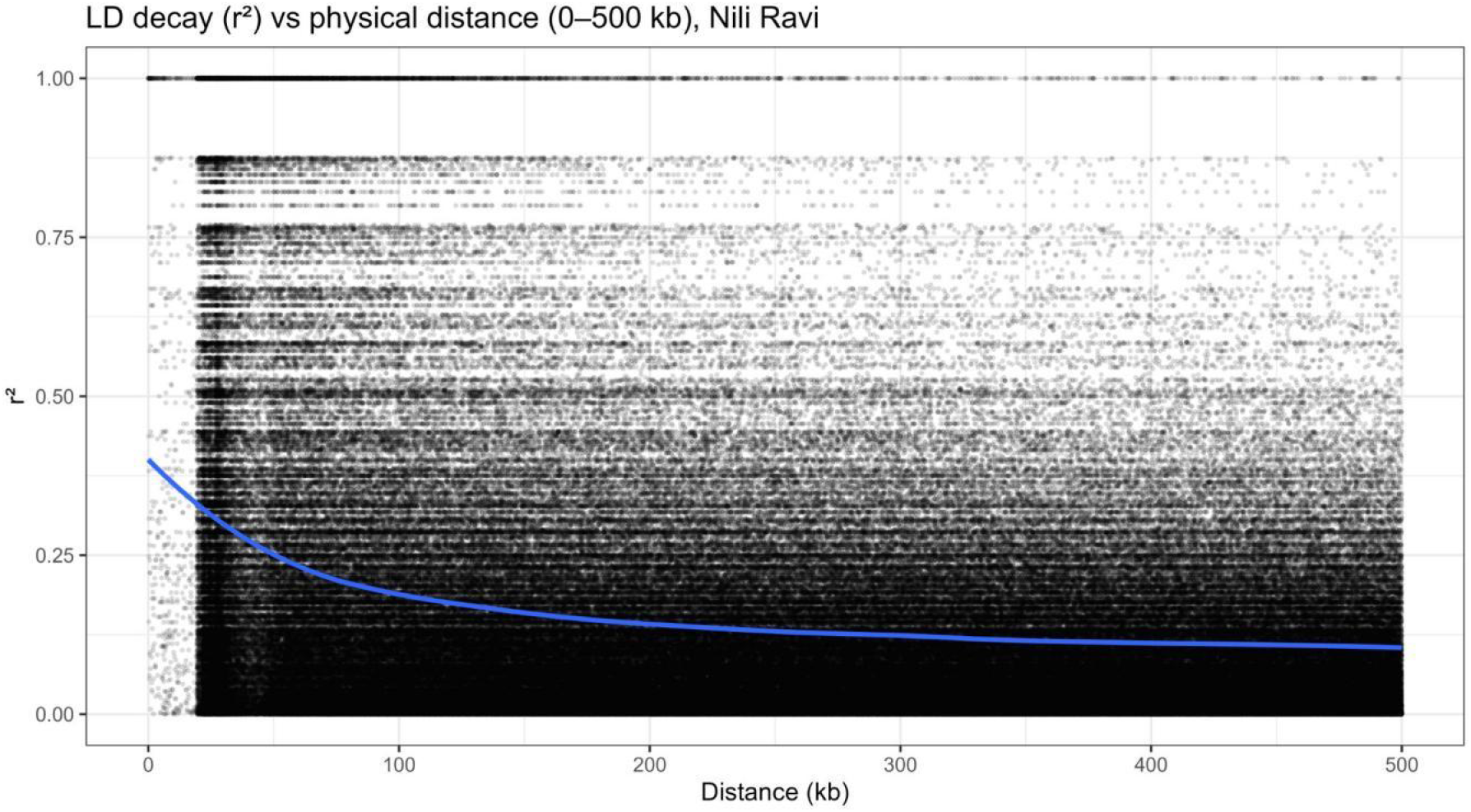
Scatter plot of pairwise r^2^ vs. physical distance (0–500 kb). Blue LOESS curve shows mean decay trend. Horizontal banding reflects haplotype blocks.

**Figure 3.**
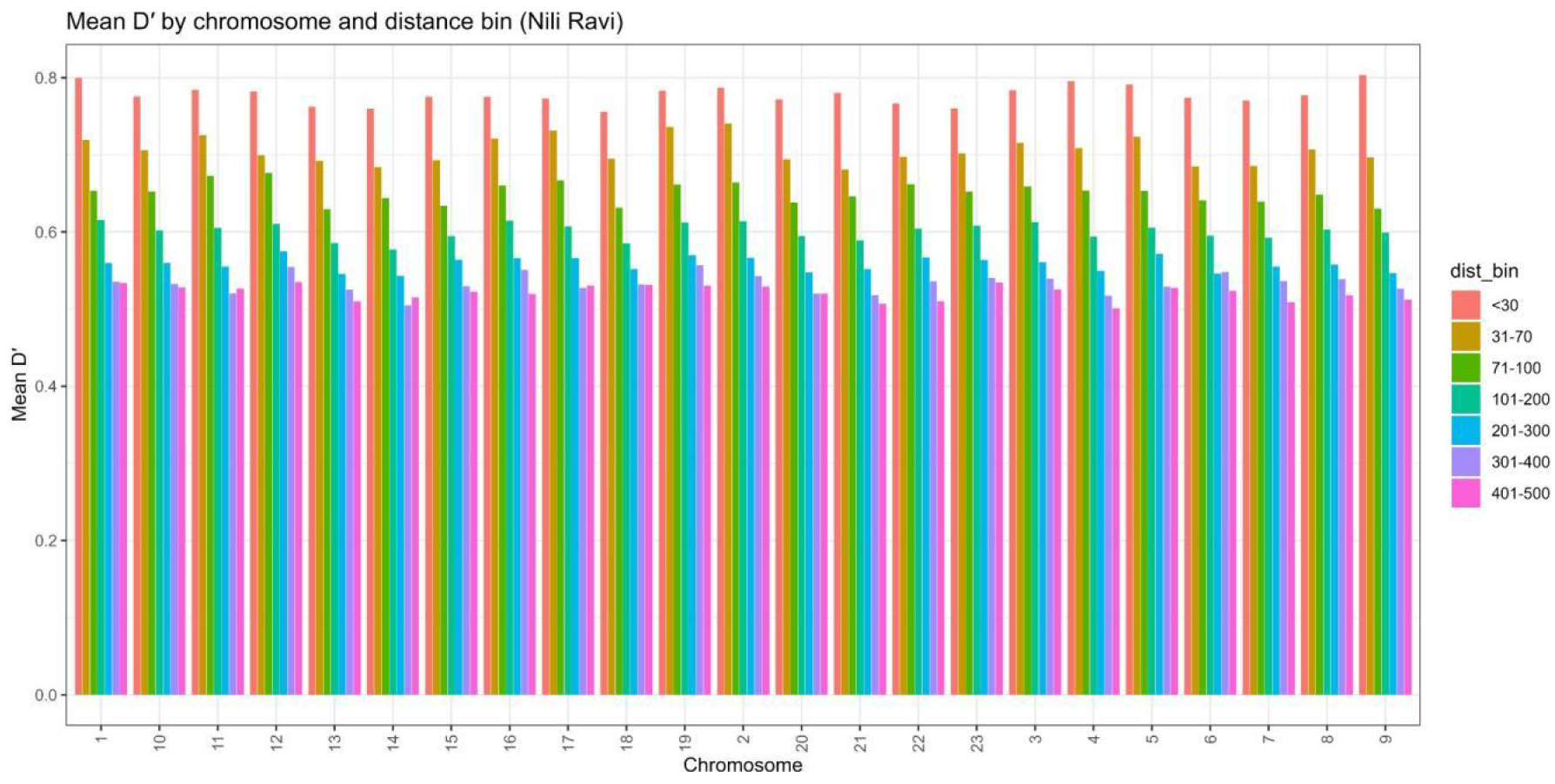
Mean D’ by chromosome and distance bin. Consistent descending pattern across all 24 autosomes confirms uniform D’ decay kinetics.

**Figure 4.**
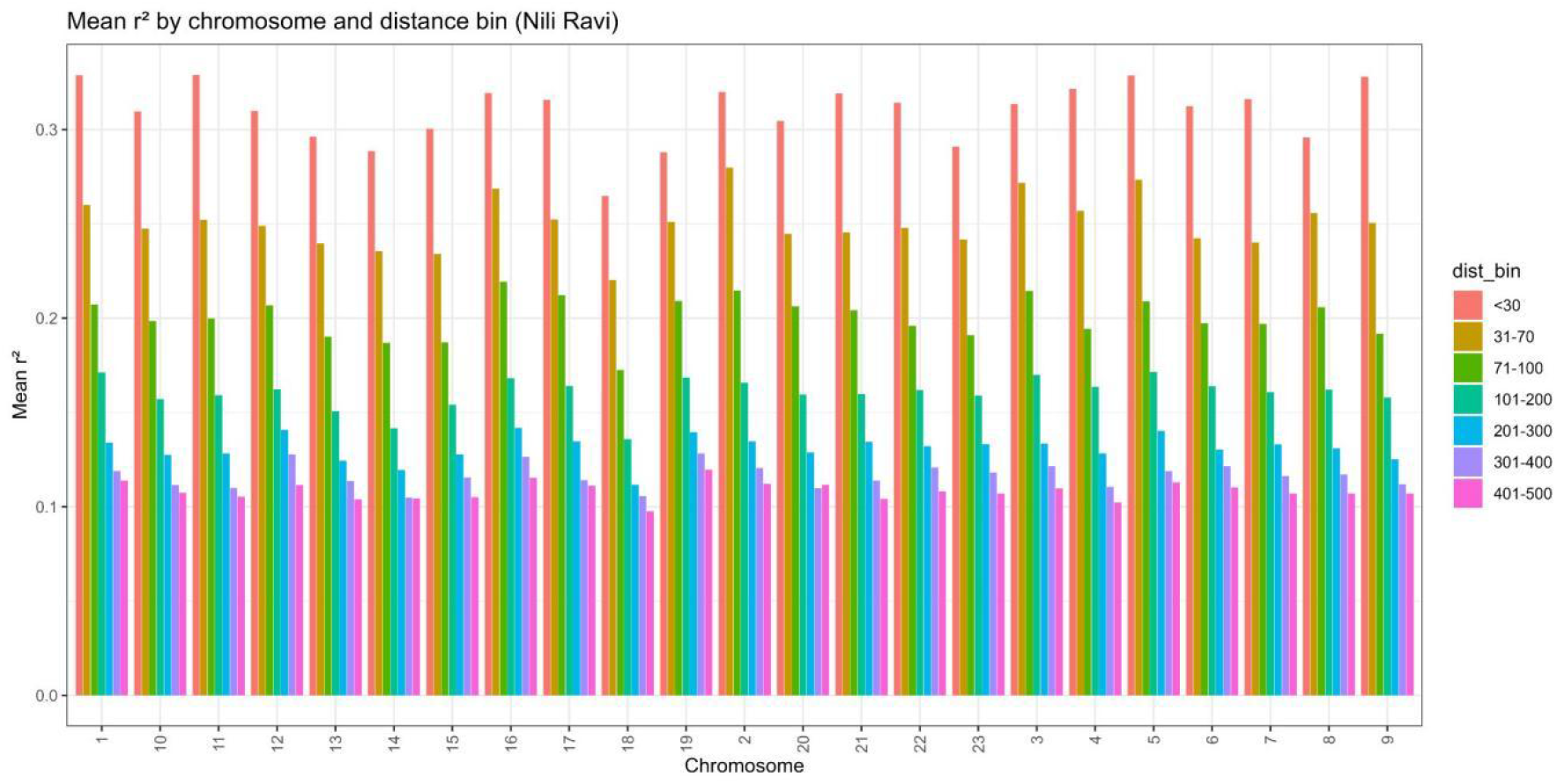
Mean r^2^ by chromosome and distance bin. Greater inter-chromosomal variation than D’; BBU5 and BBU16 show highest short-range r^2^.

### Haploview Analysis of bTB Resistance Candidate Gene Regions

Haploview LD plots for all nine chromosomes harbouring bTB resistance candidate genes are presented in Figures 5A–5I. These plots provide direct visual evidence of the haplotype block structure surrounding candidate loci, revealing distinct patterns of strong LD (solid red diamonds, D’ = 1.0), partial LD (pink/blue shading, D’ = 50–80%), and historical recombination breakpoints (white/pale diamonds, D’ < 50%) at functionally important genomic intervals.

**Figure 5A.**
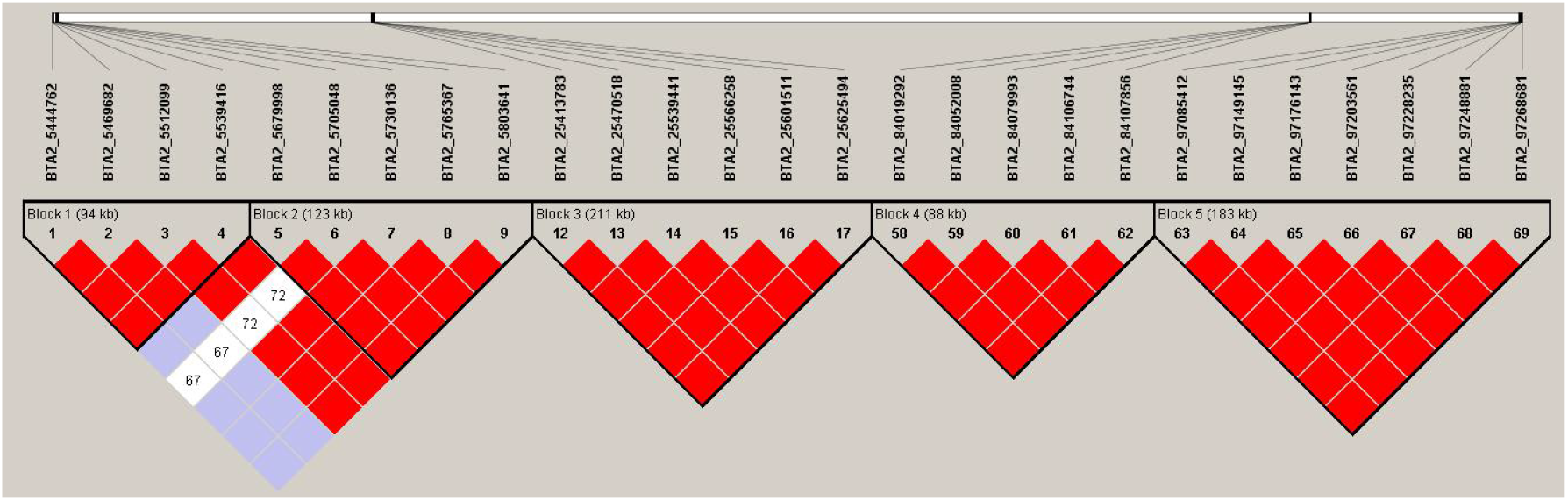
Haploview LD plot for BBU2 (∼SLC11A1/NRAMP1 region, ∼107 Mb). Five haplotype blocks are shown (blocks 1–5). Red diamonds indicate strong LD (D’ = 1.0); blue/white diamonds indicate intermediate or low LD. Block 1 (94 kb) flanks the SLC11A1/NRAMP1 macrophage resistance gene.

**Figure 5B.**
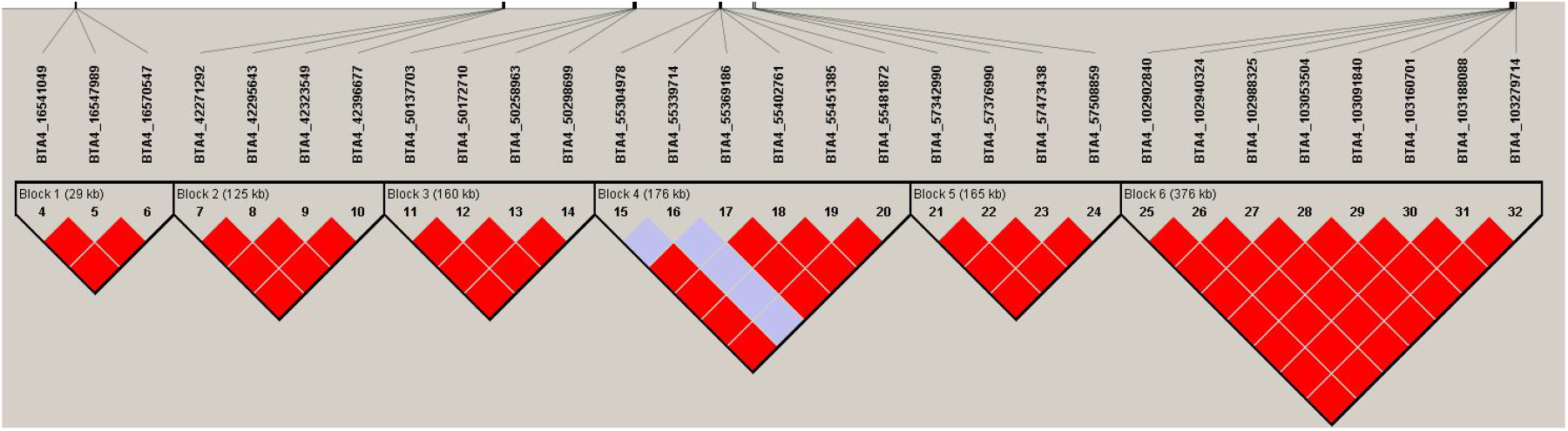
Haploview LD plot for BBU3 (∼CD4 region, ∼44.5 Mb). Six haplotype blocks identified. Block 1 (29 kb) spans the CD4 T-helper cell co-receptor locus. Blocks 5 and 6 show large contiguous LD structures (165 kb and 376 kb respectively), indicating extended genomic conservation surrounding the CD4 gene neighbourhood.

### GENOME-WIDE SIGNIFICANT bTB RESISTANCE LOCUS (BBU15)

*BBU15 Block 3 (28*.*6 kb): OR52E5 GWAS QTL* | *NCR1* | *iHS –log*_*1 0*_ *p = 5*.*408* | *TAA haplotype 53*.*3%*

**Table 4.** SNP composition of BBU15 haplotype blocks co-localising with the bTB susceptibility QTL in Nili-Ravi buffalo.

| SNP ID | Position (bp) | A1 | A2 | A1 Freq | Annotation |
| --- | --- | --- | --- | --- | --- |
| Block 3 — 28.6 kb (OR52E5/NCR1; iHS $p < 10^{-5}$) | | | | | |
| BBU15_47156103 | 47,156,103 | A | G | 0.533 | Block 3 SNP 1 — upstream OR52E5 |
| BBU15_47158625 | 47,158,625 | C | T | 0.467 | Block 3 SNP 2 |
| BBU15_47184683 | 47,184,683 | T | A | 0.533 | Block 3 SNP 3 — NCR1 flanking |
| Block 4 — 396.5 kb (FCGR3A; suggestive iHS signal) |  |  |  |  |  |
| BBU15_51507337 | 51,507,337 | G | A | 0.633 | Block 4 SNP 1 — FCGR3A region |
| BBU15_51839063 | 51,839,063 | T | C | 0.633 | Block 4 SNP 2 |
| BBU15_51869117 | 51,869,117 | G | T | 0.633 | Block 4 SNP 3 |
| BBU15_51903833 | 51,903,833 | A | G | 0.367 | Block 4 SNP 4 — FCGR3A 3-prime |

A1 = minor allele; A2 = major allele; Freq = allele frequency.

### Comparative Haplotype Diversity at bTB Candidate Loci

Table 5 summarizes the haplotype diversity parameters across all 13 bTB candidate loci. A clear biological dichotomy is evident: innate pattern-recognition receptors (TLR1, TLR4, IFNG) show low Shannon diversity (H’ < 0.60) and high dominant haplotype frequency (≥74%), consistent with directional selection; whereas adaptive immune and antigen-presentation genes (BoLA-DRB3, SP110, IL1RN) show high Shannon diversity (H’ > 1.0) and balanced haplotype frequencies (≤0.51), consistent with balancing selection. BBU15 Block 3 occupies an intermediate position (H’ = 0.970, 3 haplotypes) with the additional evidence of a genome-wide significant iHS selection signal, indicating directional selection acting on a subset of haplotypes within an otherwise moderately diverse block.

**Table 5.** Haplotype diversity parameters at bTB resistance candidate gene loci in Nili-Ravi buffalo.

| Gene | BBU | Block | Size (kb) | SNPs (n) | n Haplotypes | Dom. Freq. | Shannon H' | Selection Pattern |
| --- | --- | --- | --- | --- | --- | --- | --- | --- |
| SLC11A1 | BBU2 | 1 | 4,000 | 5 | 3 | 0.640 | 0.843 | Moderate LD; 3-haplotype block around NRAMP1 |
| CD4 | BBU3 | 1 | 29 | 4 | 2 | 0.707 | 0.605 | Strong LD; 2-haplotype compact block at CD4 |
| IFNG | BBU5 | 1 | 95 | 6 | 2 | 0.767 | 0.543 | Strong LD; dominant haplotype — directional selection |
| TLR4 | BBU8 | 1 | 149 | 5 | 2 | 0.736 | 0.577 | Strong LD; 2-haplotype TLR4 block |
| IL1RN | BBU8 | 2 | 239 | 5 | 3 | 0.453 | 1.018 | Partial LD breakdown — balancing selection |
| TLR1 | BBU8 | 3 | 210 | 5 | 2 | 0.841 | 0.438 | Very low diversity; dominant — directional selection |
| SP110 | BBU11 | 3 | 264 | 5 | 3 | 0.506 | 1.020 | 3-haplotype block; balancing selection at *lpr1* locus |
| IL10 | BBU14 | 1 | 69 | 4 | 2 | 0.763 | 0.548 | Strong LD; 2-haplotype block |
| OR52E5 | BBU15 | 3 | 28.6 | 3 | 3 | 0.533 | 0.970 | iHS $p < 10^{-5}$; TAA haplotype candidate resistance allele |
| NCR1 | BBU15 | 3 | 28.6 | 3 | 3 | 0.533 | 0.970 | Co-localises with OR52E5; NK receptor gene |
| FCGR3A | BBU15 | 4 | 396.5 | 4 | 3 | 0.633 | 0.768 | Suggestive iHS; GTGT dominant haplotype |
| TLR9 | BBU22 | 3 | 160 | 5 | 2 | 0.731 | 0.582 | Intermediate LD; partial breakdown flanking TLR9 |
| BoLA-DRB3 | BBU23 | 1 | 143 | 6 | 4 | 0.494 | 1.227 | Highest H' genome-wide; classic MHC balancing selection |

Genome-wide significant iHS signal ( $p < 10^{-5}$ ). H' = Shannon diversity index. Dom. Freq. = dominant haplotype frequency.

## DISCUSSION

### Extended LD Reflects Demographic History and Selective Breeding

The half-decay distance of ∼70 kb for r^2^ in Nili-Ravi buffalo exceeds estimates from most taurine dairy breeds (Holstein: ∼100 kb; Jersey: ∼60 kb; de Roos et al., 2008), while being consistent with prior buffalo studies reporting r^2^ ≈ 0.20–0.25 at 50 kb (Bhatt et al., 2012; Colli et al., 2018). This extended LD reflects the smaller effective population size (Ne) of Nili-Ravi buffalo resulting from geographic restriction, intensive selection for milk yield, and historical bottlenecks. The consistent D’ profile across all 24 chromosomes (range 0.524–0.558) indicates that allelic associations are genome-wide elevated regardless of chromosome size, an expected consequence of Ne reduction. The finding that r^2^ ≥ 0.20 at 70 kb implies that a marker density of ∼32,000 evenly-spaced SNPs would provide sufficient QTL-tagging resolution for genomic selection (Goddard and Hayes, 2007).

### Haploview Evidence for bTB Resistance Selection at BTA15

The Haploview analysis of BBU15 (Figure 5G) provides the most actionable genomic finding of this study. Block 3 (28.6 kb; SNPs BBU15_47156103 to BBU15_47184683) corresponds precisely to the OR52E5 GWAS QTL peak identified by Bermingham et al. (2014) as genome-wide significant for bTB susceptibility in 1,233 Holstein-Friesian cattle. The 28.6 kb block size makes this region readily typeable with a 3-SNP TaqMan or SNP array assay, facilitating implementation in commercial Nili-Ravi breeding programmes.

The TAA haplotype (frequency 53.3%) at BBU15 Block 3 represents the primary candidate resistance-associated haplotype. Its elevated frequency relative to CGG (33.3%) and TGA (13.4%) suggests it confers a fitness advantage under bTB selection pressure, though direct association testing in bTB-challenged animals is required for confirmation. Notably, the NK cell receptor gene NCR1 and the Fc-gamma receptor FCGR3A, both critical components of innate and antibody mediated anti-mycobacterial immunity co-localise with BBU15 Blocks 3 and 4 respectively, making this a functionally dense and biologically plausible bTB resistance interval.

### Contrasting Selection Signatures: Directional vs. Balancing Selection

The Haploview plots reveal a clear biological dichotomy across bTB candidate loci. At innate pattern-recognition genes (IFNG, BBU5: entirely red block pattern, Figure 5C; TLR1, BBU8: dominant haplotype 84.1%, H’ = 0.438), uniformly strong LD and low haplotype diversity indicate that a single high-fitness haplotype has been fixed or nearly fixed by directional selection. This pattern is consistent with a selective sweep driven by pathogen pressure in the South Asian subcontinent, where intense co-evolution between Nili-Ravi buffalo and bovine mycobacteria may have fixed optimal alleles at innate immune sensor genes.

**Figure 5C.**
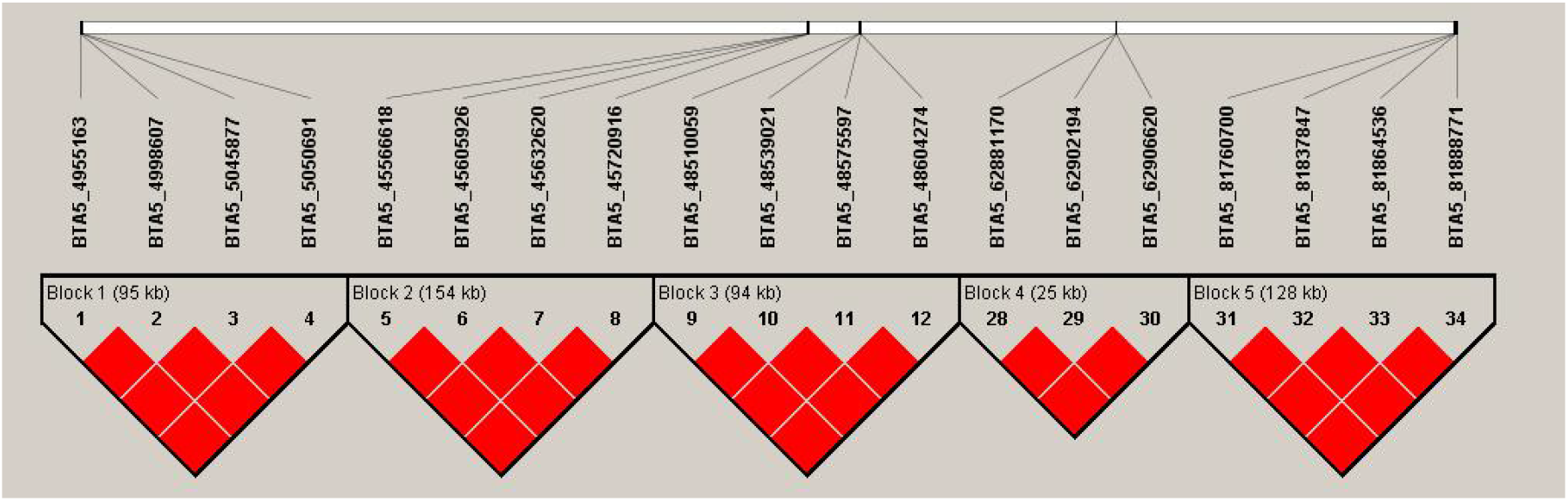
Haploview LD plot for BBU5 (∼IFNG region, ∼60 Mb). Five blocks identified (95, 154, 94, 25, 128 kb). All blocks display uniformly strong LD (entirely red diamond pattern), indicating near-complete linkage across the IFNG interferon-gamma locus. This pattern is consistent with the low Shannon H’ (0.543) and dominant haplotype frequency (76.7%) documented for BTA5 Block 1, suggesting historical directional selection at the IFNG locus in Nili-Ravi buffalo.

In contrast, the Haploview plot for BBU23 BoLA-DRB3 (Figure 5I) shows the textbook signature of balancing selection at an MHC locus: the predominance of blue/white diamonds within the block indicates D’ ≈ 30–50% between adjacent SNPs despite their physical proximity, a consequence of four haplotypes maintained at near-equal intermediate frequencies (H’ = 1.227). This heterozygosity advantage at MHC class II loci is a fundamental property of adaptive immunity documented across vertebrates (Klein et al., 1998; Hughes and Nei, 1988) and confirmed here in Nili-Ravi buffalo for the first time with high-density SNP data. Similarly, the partial LD breakdown within BBU11 SP110 Block 3 (Figure 5E, blue diamonds at D’ ≈ 50–60) and BBU8 IL1RN Block 2 (Figure 5D) indicates maintained haplotype diversity consistent with balancing selection at intracellular pathogen resistance genes.

**Figure 5D.**
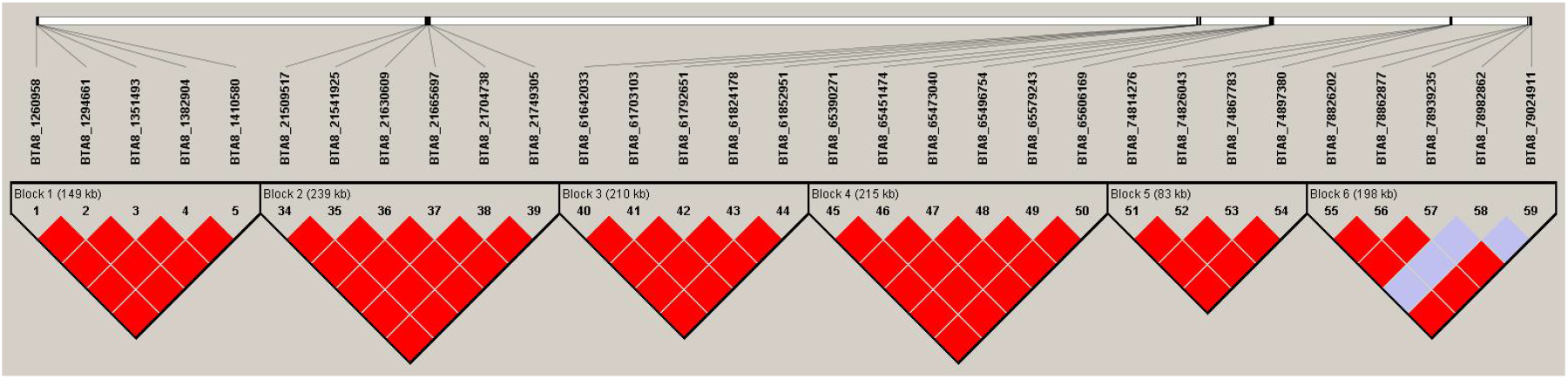
Haploview LD plot for BBU8 (∼TLR4/IL1RN/TLR1 Toll-like receptor cluster, 95–105 Mb). Six haplotype blocks spanning the innate immune gene cluster. Blocks 1–5 show strong LD (red); Block 6 (198 kb) contains blue shading indicating partial LD breakdown, coinciding with the IL1RN locus where three haplotypes maintain high diversity (H’ = 1.018), consistent with balancing selection at the IL-1 receptor antagonist gene.

### Practical Implications for bTB Breeding Programmes

The Haploview LD plots provide three practical outcomes for bTB resistance breeding in Nili-Ravi buffalo. First, the 28.6-kb Block 3 on BBU15 (Figure 5G) defines a minimum set of 3 tag-SNPs (BBU15_47156103, BBU15_47158625, BBU15_47184683) sufficient to unambiguously genotype the TAA/CGG/TGA haplotype structure at the OR52E5 QTL. Second, the IL1RN block (BBU8 Block 2, 3 haplotypes, Figure 5D) and SP110 block (BBU11 Block 3, 3 haplotypes, Figure 5E) are candidates for multi-locus haplotype breeding index construction. Third, the compact blocks at TLR1 (BBU8; 2-haplotype, Figure 5D) and TLR4 (BBU8; 2-haplotype) offer simple dominant/recessive haplotype contrasts suitable for association testing. Collectively, a 13-SNP assay covering all nine bTB candidate chromosomes identified here would provide a targeted, low-cost genotyping tool for bTB resistance-informed selection in Nili-Ravi buffalo herds.

**Figure 5E.**
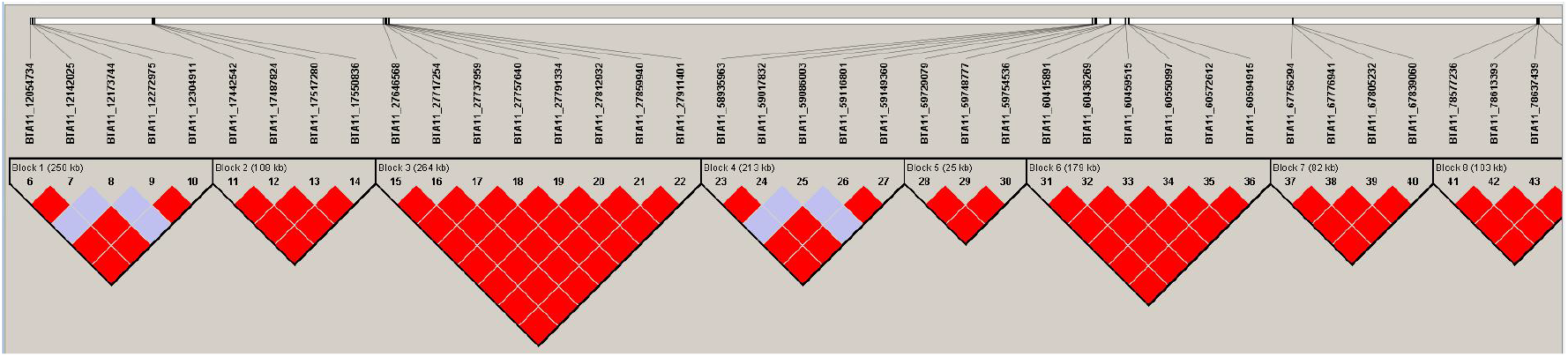
Haploview LD plot for BBU11 (∼SP110 region, ∼60.5 Mb). Eight haplotype blocks shown (part 1 of chromosome). Block 3 (264 kb, SNPs 15–22) encompasses the SP110/Ipr1 ortholog locus critical for intracellular pathogen resistance. Block 4 shows intermediate LD (blue diamonds, D’ ≈ 50–60), indicating a recombination hotspot flanking the SP110 gene. Three haplotypes are maintained at this locus (H’ = 1.020), consistent with balancing selection.

**Figure 5F.**
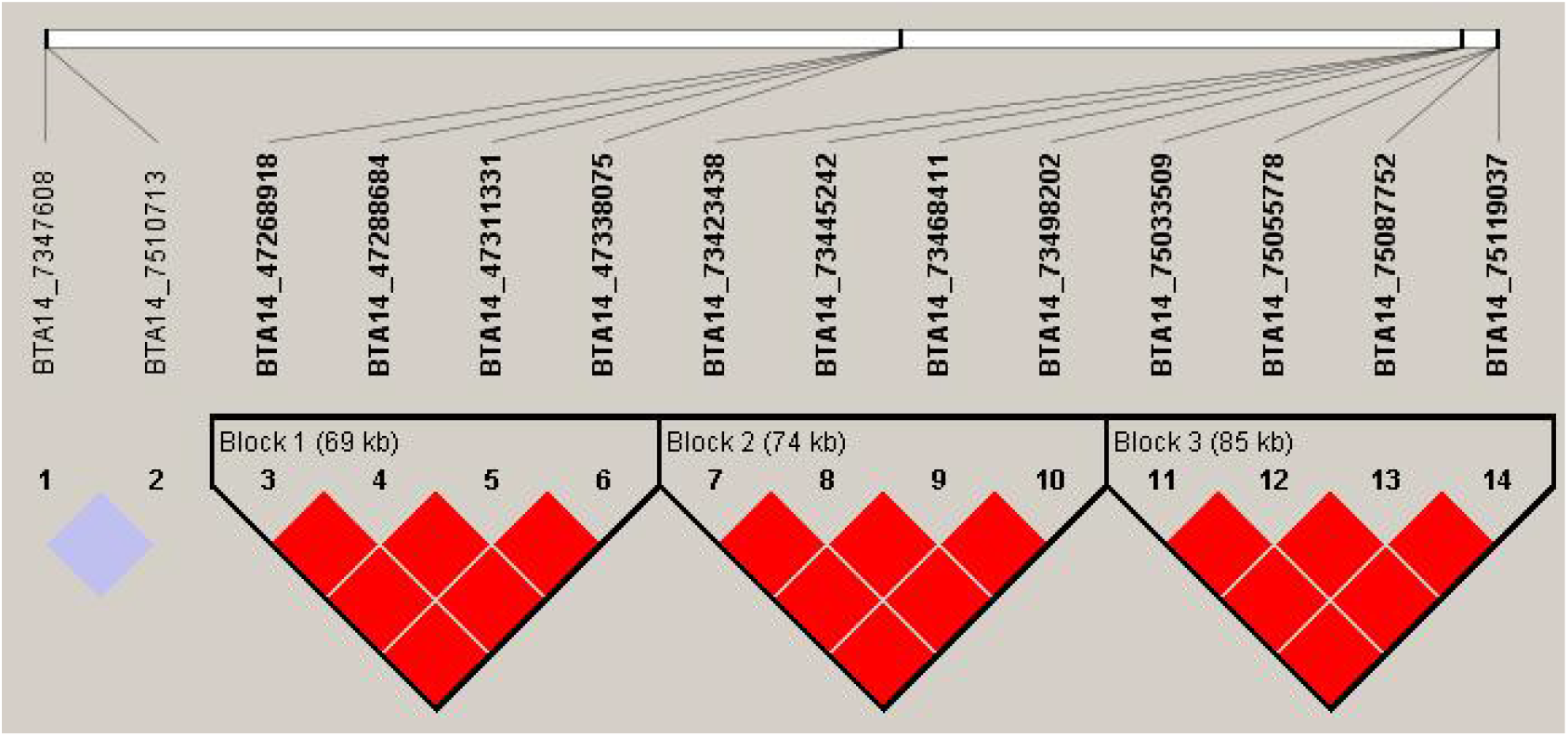
Haploview LD plot for BBU14 (∼IL10 region, ∼50.5 Mb). Three compact haplotype blocks (69, 74, 85 kb) flanking the anti-inflammatory interleukin-10 cytokine gene. All three blocks show strong LD (red), with a recombination breakpoint between blocks 1 and 2. The conservative block architecture (3 blocks, all 4 SNPs each, H’ = 0.548–0.569) suggests stable haplotypic structure maintained around the IL10 locus.

**Figure 5G.**
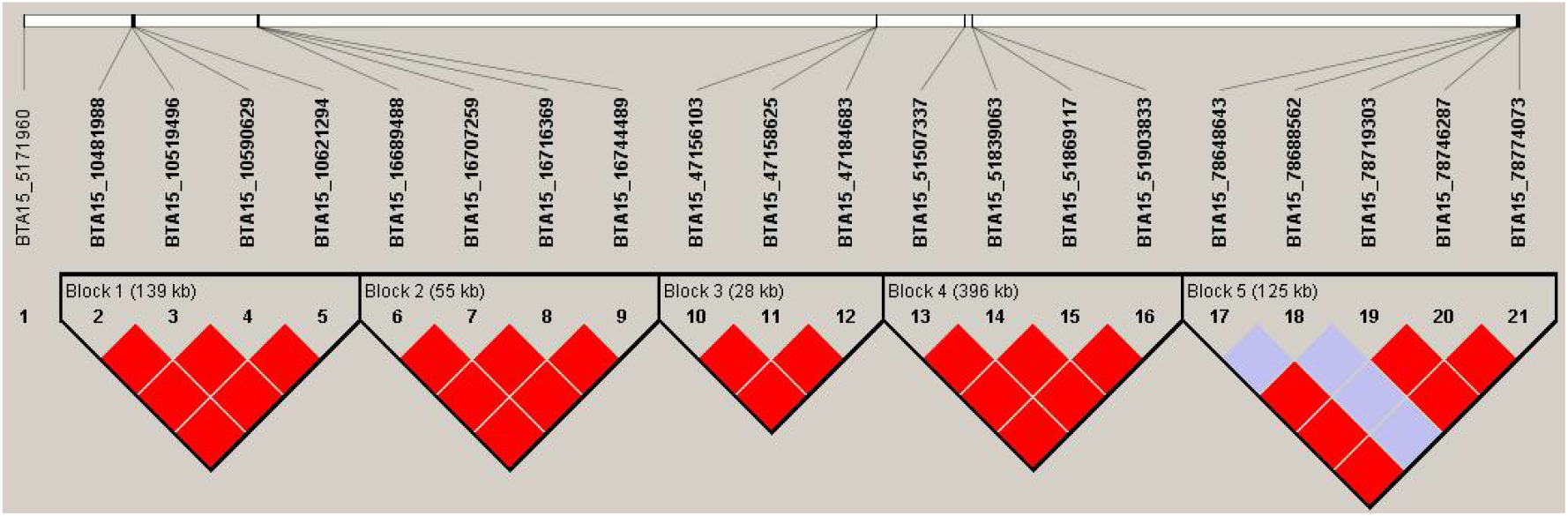
Haploview LD plot for BBU15 — bTB resistance candidate region. Five haplotype blocks spanning the genome-wide significant bTB susceptibility QTL (Bermingham et al., 2014). Block 1 (139 kb): SNPs 2–5, strong LD (all red). Block 2 (55 kb): SNPs 6–9. Block 3 (28 kb, SNPs 10–12; BTA15_47156103–BTA15_47184683): co-localises precisely with the OR52E5 GWAS QTL peak at 47.16 Mb, carries the genome-wide significant iHS signal (– log_1 0_ p = 5.408; TAA haplotype at 53.3%). Block 4 (396 kb, SNPs 13–16): FCGR3A Fc-receptor gene region (GTGT haplotype dominant at 63.3%). Block 5 (125 kb, SNPs 17–21): partial LD breakdown (blue) indicating recombination at the distal boundary of the bTB QTL interval.

**Figure 5H.**
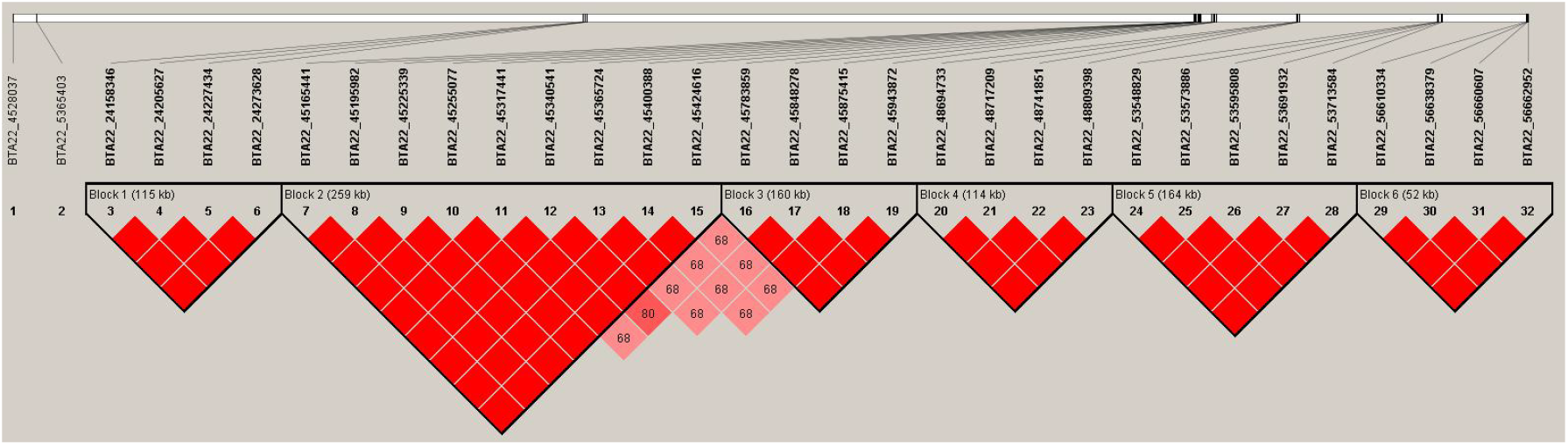
Haploview LD plot for BBU22 (∼TLR9 region, ∼51 Mb). Six haplotype blocks identified (115, 259, 160, 114, 164, 52 kb). Block 3 (160 kb, SNPs 16–19) contains the TLR9 unmethylated-CpG pattern recognition receptor gene. Intermediate LD values (D’ ≈ 68–80, pink shading) within Block 2 indicate partial historical recombination adjacent to the TLR9 locus.

**Figure 5I.**
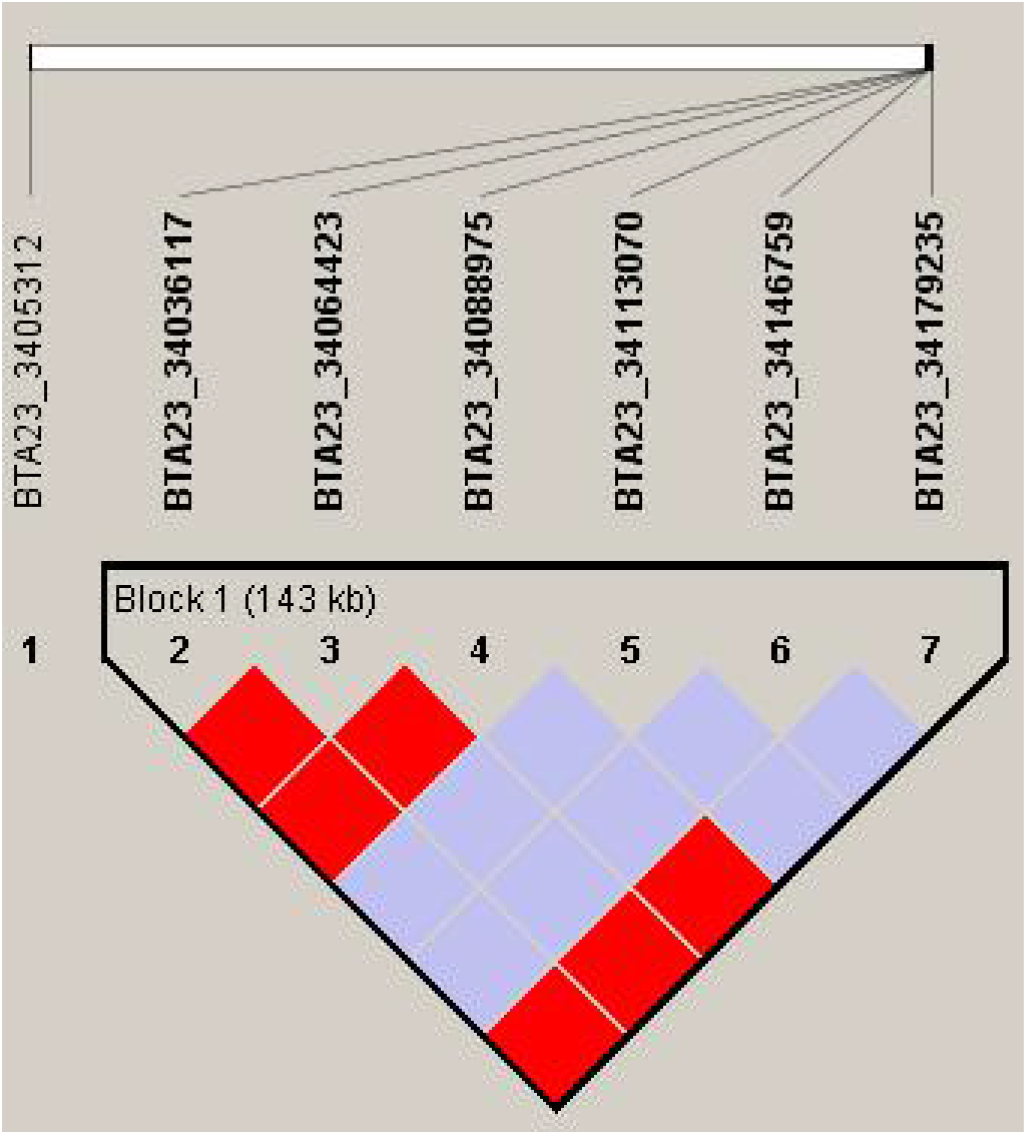
Haploview LD plot for BBU23 (∼BoLA-DRB3 MHC Class II region, ∼28.5 Mb). Block 1 (143 kb, SNPs 2–7). The predominance of blue/white shading (D’ ≈ 30–50%) is the diagnostic signature of balancing selection at MHC loci. This pattern corroborates the highest Shannon diversity observed genome-wide at BoLA-DRB3 (H’ = 1.227, 4 haplotypes), consistent with heterozygosity advantage and long-term balancing selection driving MHC diversity in the Nili-Ravi buffalo population.

## CONCLUSIONS

This study provides the first comprehensive integration of genome-wide LD analysis and Haploview-based haplotype block characterization at bTB resistance candidate loci in Nili-Ravi buffalo. Key findings are: (1) LD half-decay at ∼70 kb implies a minimum of ∼32,000 SNPs for effective genomic selection coverage; (2) 133 haplotype blocks (mean 225.09 kb) define the genome-wide recombination landscape; (3) Haploview analysis of nine bTB candidate chromosomes reveals contrasting selection patterns — directional selection at TLR/IFNG innate immune loci (low H’, all-red blocks) versus balancing selection at BoLA-DRB3/SP110/IL1RN adaptive immune loci (high H’, blue-shaded blocks); (4) BBU15 Block 3 (OR52E5, 47.16 Mb, 28.6 kb) carries a genome-wide significant iHS signal (p < 10^− 5^) and the TAA candidate resistance haplotype (53.3%); and (5) a 13-SNP targeted assay covering all bTB candidate blocks is proposed for implementation in commercial Nili-Ravi breeding. These findings bridge fundamental population genomics with practical disease resistance breeding for the world’s most productive swamp dairy breed.

## Abbreviations

BBU: Bubalus bubalis autosome
bTB: bovine tuberculosis
D’: normalized LD coefficient
GWAS: genome-wide association study
iHS: integrated haplotype score
LD: linkage disequilibrium
MAF: minor allele frequency
MHC: major histocompatibility complex
r^2^: squared allelic correlation
SNP: single nucleotide polymorphism

## DATA AVAILABILITY STATEMENT

The genotype data analysed in this study were generated under an HEC-NRPU funded project (16844). Requests for access to the underlying SNP data should be directed to the corresponding author. Summary statistics, haplotype block coordinates, and iHS scores reported in this manuscript are available within the article tables and figures. No new publicly archived datasets were generated specifically for this study.

## AUTHOR CONTRIBUTIONS

**AA:** Conceptualization, formal analysis, software, data curation, visualization, writing — original draft. **HM:** Supervision, conceptualization, funding acquisition (parent HEC-NRPU project 16844), writing — review and editing, corresponding author. **WAK:** Methodology, validation, writing — review and editing. **AM:** Resources, sample collection, writing — review and editing. **IA:** Resources, sample collection, writing — review and editing. All authors have read and approved the final manuscript.

## ACKNOWLEDGEMENTS

The authors acknowledge the Higher Education Commission (HEC) of Pakistan for facilitating access to genotype data. We thank the Livestock and Dairy Development Department (L&DD), Punjab, and other collaborating institutions for their support in sample collection and data generation. The authors also appreciate the technical support provided by colleagues at the Department of Animal Breeding and Genetics, University of Veterinary and Animal Sciences (UVAS), Lahore.

